# The microbiota affects stem cell decision making in *Hydra*

**DOI:** 10.1101/2024.08.20.608462

**Authors:** Jinru He, Alexander Klimovich, Sabine Kock, Linus Dahmke, Sören Franzenburg, Thomas C.G. Bosch

**Affiliations:** Zoological Institute, University of Kiel, Kiel, Germany; Institute of Clinical Molecular Biology, University of Kiel, Kiel, Germany

**Keywords:** host-microbe interactions, phylosymbiosis, pattern formation, cnidaria

## Abstract

Research on microbial communities colonizing animals has revealed that the microbiota, despite its typical containment to surfaces, influences virtually all organ systems of the host. In absence of a natural microbiota, the host’s development can be disturbed, but how developmental programs are affected by the microbiota is still poorly understood. Removing the microbiota from *Hydra*, a classic model animal in developmental biology, causes drastic developmental malformations and leads to polyps that temporarily lack the ability to bud. Recolonizing non-budding germfree *Hydra* with bacteria reverses this budding inhibition. Single-nucleus ATAC-seq detected a unique chromatin landscape associated with the non-budding phenotype. Single-cell RNA-seq and trajectory-based differential expression analysis showed that epithelial stem cell decision making is disturbed in non-budding polyps, whereby key developmental regulators are not expressed. This process is reversible by adding back bacteria. Transcriptionally silencing of one of the genes that failed to be activated in non-budding animals, GAPR1, led to polyps that have a significantly reduced budding capacity. The results show that maintaining a species-specific microbiota may enable the animal host to maintain its developmental program.

**Significance Statement:** Animal developmental programs work within the context of coevolved associations with microbes. Here, we provide mechanistic evidence of the involvement of the microbiota in maintaining the pattern formation program of *Hydra* with the asexual formation of buds in the lower part of the body column. We demonstrate that in the absence of bacteria both the epigenetic and transcriptomic landscape is changed and that key regulatory factors are not expressed, causing changes in stem cell trajectories that result in loss of budding capacity. This study provides a new perspective on the role that microbiota plays during animal development and evolution.

**One Sentence Summary:** Microbiota interfere with *Hydra*’s asexual reproduction via modulating its stem cell differentiation programs.

## Introduction

Multicellular organisms including humans form lifelong associations with microbial communities that can contain archaea, bacteria, fungi and viruses (Bosch and McFall-Ngai 2011; McFall-Ngai et al 2013, Dominguez-Bello et al 2019). As research on microbial communities colonizing animals and plants progressed over the past decades, it revealed that these microbiota, despite their containment to surfaces, influence the development and function of virtually all organ systems of an animal host, resulting in both local and systemic effects (Hill and Round 2021). The interactions between animals and their bacteria are probably as ancient as animals themselves (Rawls et al 2006) and have profoundly impacted the evolution of both, the hosts and their microbes. Phylosymbiosis describes the links between evolutionary relationships of host species and their associated microbiota (Lim and Bordenstein 2020). Identified linkages indicate that the host genotype may select for certain microbes (Lynch and Hsiao, 2019). The terms “holobiont” and “metaorganism” were introduced to collectively describe the host organism and all its associated microorganisms (Theis et al 2016; Roughgarden et al 2018). In this context, Gould and Lewontin were far ahead of their time, when they commented “… *organisms must be analyzed as integrated wholes*,” (Gould and Lewontin, 1979). Changes in the composition or abundance of the microbiota of a host can result in developmental disorders, and in humans some have been associated with chronic diseases (von Frieling et al 2018; Barcik et al 2020; de Vos et al 2022). However, the mechanisms by which bacteria influence the development of their hosts are still largely unknown.

Evolutionary developmental biology and metaorganism research is typically studied in model organisms, of which the cnidarian *Hydra* is a prime example (Trembley, 1744; Browne 1909; Bosch 2013, Tomczyk et al 2015; Vogg et al 2019; Douglas 2019; McFall-Ngai and Bosch 2021; Holstein, 2023, Kovacevic et al 2024). Cnidaria occupy a sister phylogenetic position to bilaterians (**Fig. 1A**). *Hydra* polyps have a clearly structured oral-aboral body axis, with a head, tentacles and a foot. The body axis is established and maintained by position-dependent gene expression in which a number of cell-cell signaling pathways are involved (**Fig. 1B**). Comparative genomics has demonstrated similarities between *Hydra*’s molecular developmental toolkit and that of more complex animals (Kortschak et al 2003; Technau et al 2005; Augustin et al 2006; Erwin 2009; Wenger & Galliot 2013, Holzem et al 2024), for instance the critical role of the Wnt signaling pathway (Holstein, 2022). *Hydrá*s body wall consists of two epithelial layers, an ectoderm and an endoderm, which are separated by a mesoglea, an extracellular matrix (**Fig. 1C**). The glycocalyx covering the ectoderm is colonized by the microbiota. The ectoderm regulates body morphology, while the endoderm controls body size, as was early demonstrated by classical experiments with chimeric strains (Wanek and Campbell 1982). Stem cells of the interstitial cell lineage give rise to nerve cells, nematocytes, gland cells, and germ cells (**Fig. 1C**). Both ectodermal and endodermal epithelial cells continuously divide and move along the body axis toward the ends, where they are removed by sloughing. Well-fed *Hydra* reproduce asexually by the formation of buds (**Fig. 1B**), which develop into a fully formed individual that eventually detaches from the parental polyp. This process takes about 3.5 days and is critical for the fitness of the animal. *Hydra* experimentally depleted of all cell types except the epithelial cells are able to bud (Campbell 1976), indicating that the developmental mechanics of bud formation must depend on the ability of epithelial cells to migrate, rear-range, and adhere to one another (Otto and Campbell, 1977a). Buds are formed by tissue evagination, generating a new body axis (Bode, 2009) controlled by the Wnt/β-catenin pathway (Hobmayer et al 2000; Broun et al 2005; Philipp et al 2009; Gee et al 2010, Nakamura et al 2011). The size of an adult animal is determined by the balance between cell production and cell loss via sloughing, and bud formation. All three stem cell lineages (endodermal, ectodermal and interstitial cells) are able to divide indefinitely, enabling everlasting asexual growth (Bosch 2009; He and Bosch 2022). Single-cell RNA sequencing has been performed on a wide range of *Hydra* cell types and their differentiation states, which provided first insights into putative regulatory modules that drive cell differentiation and gene expression (Siebert et al 2020).

**Fig. 1.**
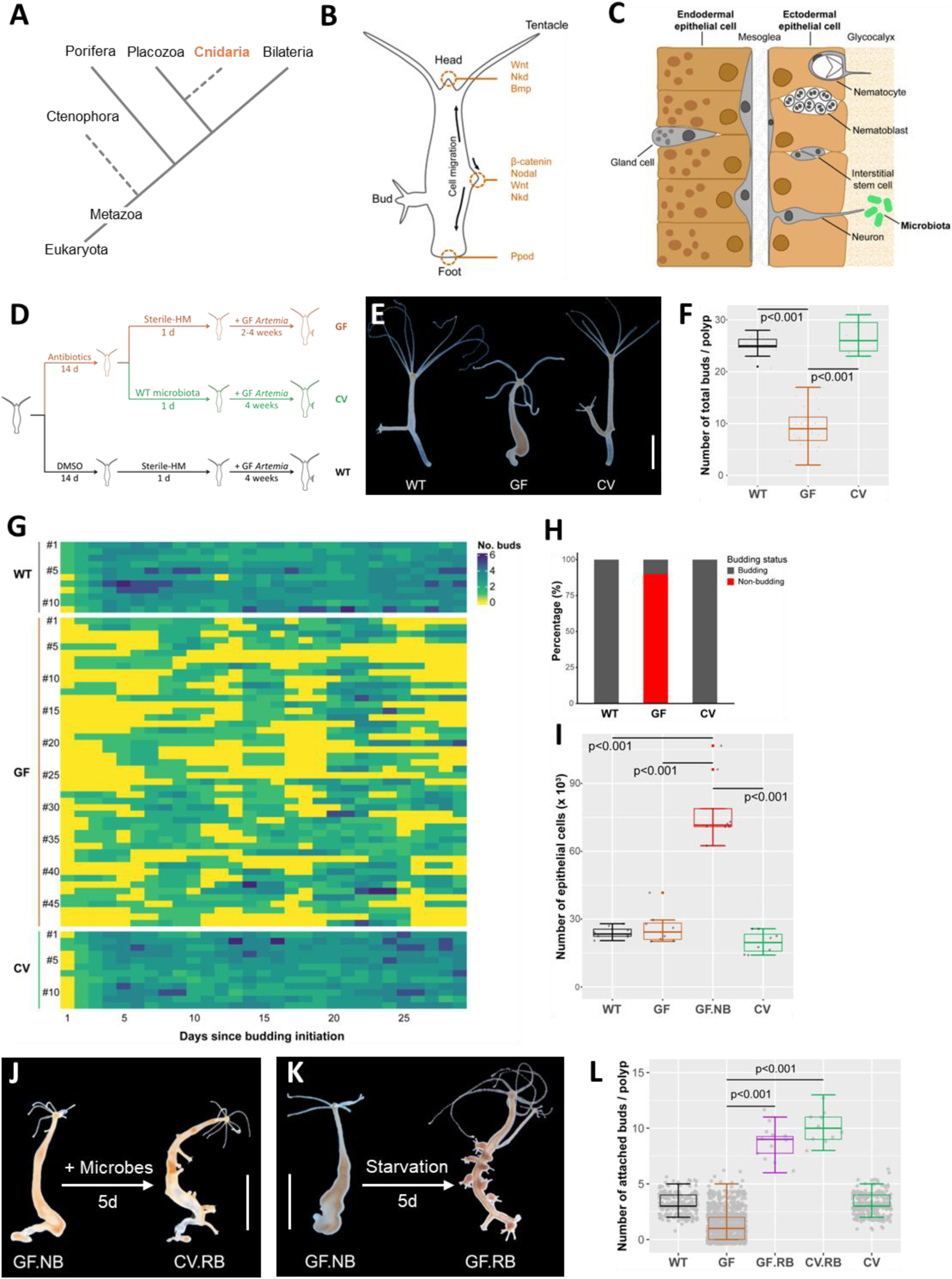
Budding is reversibly inhibited in the absence of microbes. **A**. The model organism *Hydra* is a member of the phylum Cnidaria that represents a sister group to bilaterians. **B**. The body plan of *Hydra* contains a head and tentacles at the oral end and a foot at the aboral end of the polyp. Stem cells reside in the body column and migrate towards the ends. Factors involved in developmental processes are indicated, and a bud is forming to the left of the body. **C**. The body wall of *Hydra* contains two cellular layers, the ectoderm and the endoderm, both consisting of epithelial cells. The ectoderm is covered by a glycocalyx on the outside in which the microbiota resides. A third cell lineage of interstitial cells is located within these two epithelial layers and these cells can differentiate into neurons, gland cells, nematocytes, and germline cells. **D**. Experimental setup of the standard procedure. Germ-free (GF) *Hydra* polyps were created by incubating wildtype (WT) polyps with antibiotics for two weeks. Conventionalized (CV) polyps were recolonized with native bacteria. All animals were daily fed with germ-free Artemia. **E.** GF polyps gradually stopped budding (polyps shown at day 30). Scale bar: 3 mm. **F.** The number of buds per animal was significantly lower for GF polyps compared with WT and CV. **G.** Once budding started, individual WT and CV polyps continuously produced new buds, but GF polyps frequently and repeatedly stopped budding, to then resume budding autonomously. **H.** In 4 weeks, budding inhibition occurred in 89% of GF *Hydra* (n=24), but not in WT or CV. **I.** 7-day non-budding GF *Hydra* (GF.NB) had a three-fold increased body size compared with regularly budding GF, WT, or CV polyps, as determined by epithelial cell counts. **J.** Recolonizing GF.NB animals with microbiota from WT polyps induced budding bursts (giving CV.RB), with buds emerging around the same time and growing almost along the entire body column. Scale bar: 5mm. **K.** Budding bursts also occurred in starved GF.NB polyps without exposing them to microbes (GF.RB). Scale bar: 5mm. **L**. Budding bursts in GF.RB and CV.RB resulted in a three-fold increase in budding numbers compared with regularly budding GF, WT and CV polyps.

The natural species-specific microbiota occupying the glycocalyx of *Hydra* (**Fig. 1C**) has a low complexity and is essential for development and health (Fraune and Bosch 2007; Franzenburg et al 2013). A disturbed or reduced microbiota influences the animal’s immune system (Fraune et al 2015), behavior (Murillo-Rincon et al 2017; Giez et al 2023) and development (Taubenheim et al 2020). The impact of the microbiota on the developmental program of *Hydra* is of particular interest with regard to its ability to reproduce asexually by budding. It had early been established that bud induction depends on the size of the parental polyp and its epithelial growth rate (Otto and Campbell 1977b). Five years later, it was described that some bacteria-free *Hydra* polyps were unable to form buds (Rahat and Dimentman, 1982), and involvement of a missing budding factor was proposed that might be provided exogenously by either nonsterile food or by bacteria; however, such a factor has so far not been identified. Here, we explored the impact of the microbiota on budding capacity in an attempt to identify the underlying mechanisms of this intricate host-microbiota interaction.

## Results

The impact of the microbiota on the budding capacity of *Hydra* was investigated using the genetically characterized species *Hydra vulgaris* AEP, which can be cultivated under germ-free conditions of over long periods of time. ATAC-seq (Assay for Transposase-Accessible Chromatin using sequencing) was used to assess genome-wide chromatin accessibility under germ-free conditions. Single-cell RNA sequencing (scRNA-seq) was applied to capture the molecular progression of cells under bacteria-free conditions over time, and reverse genetics was used to functionally investigate the impact of selected *Hydra* genes that were no longer expressed in non-budding germ-free animals.

### Removal of bacteria leads to emergence of *Hydra* that can no longer bud

The importance of a microbiota for developmental processes was demonstrated with germfree (GF) *Hydra* polyps that were compared with their wild-type (WT) counterparts, as previously described (Franzenburg et al 2013). In addition, to exclude the potential impacts of the antibiotics treatment used to generate GF, conventionalized (CV) polyps were created by recolonizing GF polyps with supernatant from homogenized WT *Hydra* (Materials and Methods). The standard experimental setup is shown in **Fig. 1D**. Once budding started, WT and CV *Hydra* continued to produce new buds, but GF polyps frequently entered a non-budding state (**Fig. 1E, G**). Tracking individual animals over a period of four weeks revealed that the total number of buds per polyp was significantly lower for GF polyps than for controls **(Fig. 1F**). After an extended non-budding phase, individual GF animals could spontaneously resume budding for a short time and re-enter the non-budding state again (**Fig. 1G**). Overall, non-budding occurred in 89% of GF polyps but was virtually absent in WT or CV (**Fig. 1H**). Epithelial cell counts confirmed that non-budding GF animals grew larger than budding WT and CV controls, pointing to a disturbed stem cell activity. Since the body size of budding GF animals remained normal (**Fig. 1I**), we therefore distinguished between budding GF individuals and non-budding GF individuals (referred to as “GF.NB”).

Although some GF animals could spontaneously escape the budding arrest for a short time, before returning into a non-budding state, re-colonization of GF.NB polyps with microbiota from WT polyps effectively induced/reverted the polyps to a stable budding state (**Fig. 1G**, lower panel, **Fig. 1J**). The resulting buds in these recolonized animals developed into normal polyps, as long as microbes were present. Interestingly, the same re-budding phenotype was also observed when GF.NB polyps were starved for 5 days (**Fig. 1K**), indicating that the recolonization somehow interfered with the host’s metabolism. Apparently, under regular feeding conditions a budding inhibitor accumulated in germ-free animals. These findings suggest that microbes have a drastic influence on *Hydra* developmental processes, affecting the pattern formation system in adult polyps and influencing the ability to form buds and to proliferate asexually.

### Germ-free non-budding polyps show an altered gene expression profile in both ectodermal and endodermal epithelial stem cells

To explore how bacteria influence the development program and which processes are affected, transcriptional changes were characterized by scRNAseq in budding control (WT and CV) and germfree (GF) polyps, and in non-budding germ-free (GF.NB) polyps. Sequences from single-cell libraries were integrated with available scRNAseq data from polyps cultured under normal conditions (Siebert et al., 2019), to produce a reference *Hydra* cell atlas using Uniform Manifold Approximation and Projection (UMAP) (**Fig. 2A**). The integrated atlas comprised 80,888 high-quality single-cell transcriptomes from the different samples. The integrated projection recovered the molecular signatures of the three stem cell lineages of *Hydra*, with trajectories for each lineage, and reflected the differentiation paths from stem cells to terminally differentiated cell types (**Fig. 2A**). Since epithelial cells, rather than interstitial cells or nerve cells, are the primary determinant of most, if not all, of *Hydra*’s developmental characters (Sugiyama and Fujisawa 1978), we focused on the profiles of ectodermal and endodermal epithelial cells. A comparison of cell composition between GF.NB, WT, GF and CV polyps revealed a substantial accumulation of cells with a molecular profile resembling that of the foot epithelial cells (**Fig. 2B**, arrowheads), indicating that stem cell decision making is disturbed in GF.NB polyps. To characterize the epithelial stem cells of the body column in GF.NB polyps, scRNA-seq was performed on GF and GF.NB polyps from which the head and foot had been removed (**Fig. 2C**). As expected, in regularly budding GF polyps terminally differentiated head or foot cells could no longer be detected. In contrast, in the body column of non-budding polyps we uncovered in both the ectodermal and the endodermal epithelial lineages a population of terminally differentiated foot cells (arrowheads in **Fig. 2C**) in addition to head-like endodermal and ectodermal cells. These observations suggest that the budding inhibition induced by the absence of the microbiome is accompanied by the appearance of foot-specific epithelial cells in the gastral region. We next tested whether the absence of bud formation is associated with the expression of foot-specific genes and the production of foot-specific proteins in the body column of GF.NB polyps. For this, in-situ hybridization of the foot marker gene L-rhamnose-binding lectin CSL3-like (RBL) was used (Wenger et al 2014). This marker was exclusively expressed in the foot region of regularly budding GF, but ectopic expression was observed well outside the foot region up to the area around the head of GF.NB (arrowheads in **Fig. 2D**). In *Hydra*, differentiated ectodermal cells of the foot region contain a peroxidase activity encoded by the *ppod1* gene, and this can also be used as a marker for foot-specific differentiation processes (Hoffmeister-Ullerich et al 2002). Staining for this foot-specific peroxidase activity confirmed that ectopic terminally differentiated epithelial cells accumulated in the body column of GF.NB (**Fig. 2E-F**). Taken together, the data revealed that in GF.NB polyps undifferentiated epithelial stem cells accumulated in the body column, and these cells can ectopically differentiate into foot cells, but not head cells. Thus, the absence of colonizing bacteria seems to affect epithelial stem cell decision making and to cause a temporary loss of self-regulation to maintain tissue homeostasis and budding.

**Fig. 2.**
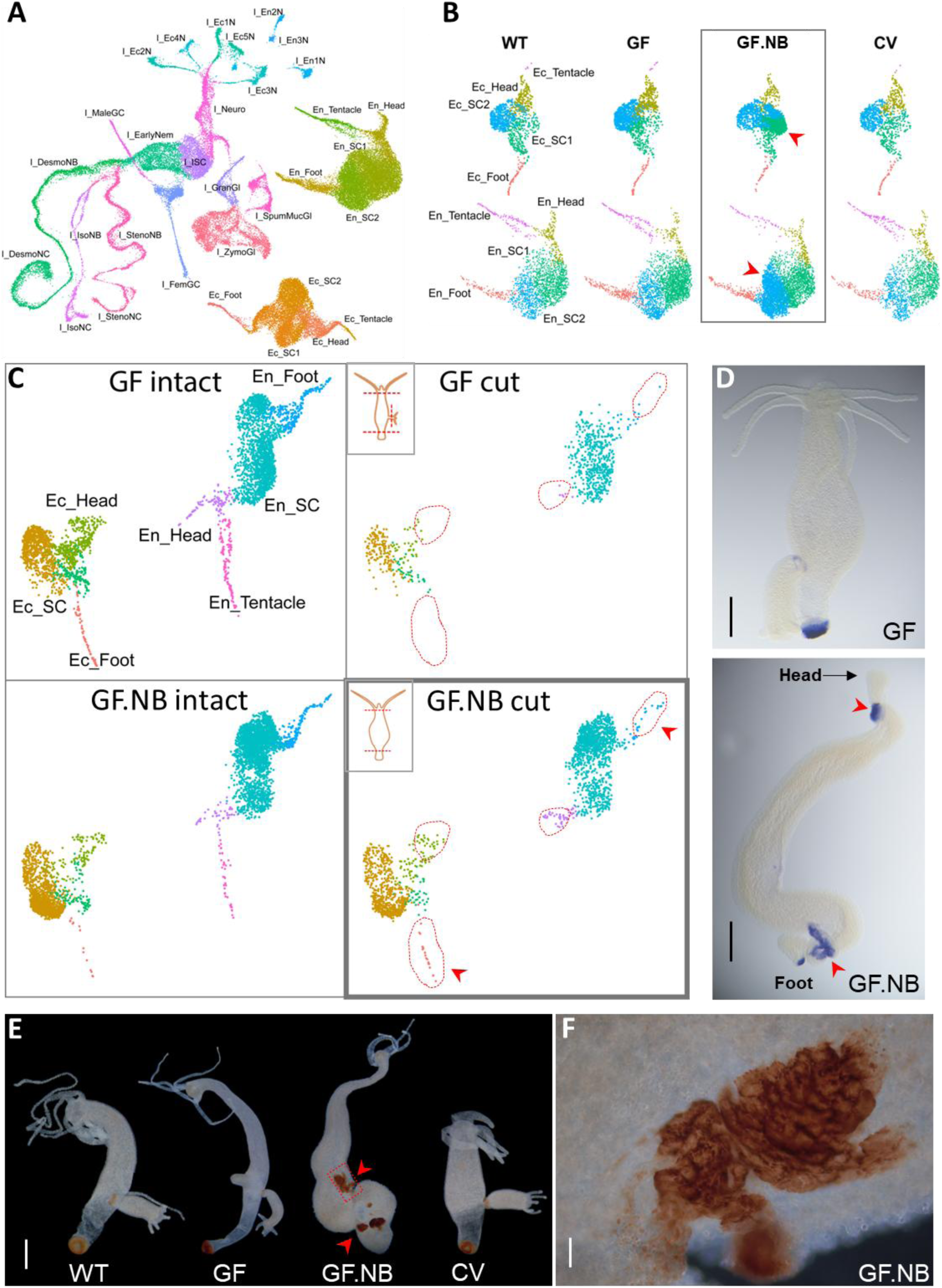
Single cell transcriptome analysis reveals redirected epithelial stem cell differentiation. **A**. Uniform Manifold Approximation and Projection (UMAP) representation of integrated single-cell transcriptomes. The projection recuperates the three cell lineages (En: endodermal; Ec: ectodermal; I: interstitial) of *Hydra*, visualizing well-preserved differentiation paths for each lineage (designated by color) to terminally differentiated cell types. **B**. Comparison of epithelial cell composition between sample groups, with 2 animals per group. In GF.NB, both ectodermal and endodermal epithelial stem cell subpopulations accumulated whose transcriptional profile demonstrated closer similarity to the foot cell identity (red arrows). **C**. Experimental removal of the heads and foots did not deplete all terminally differentiated epithelial cells in GF.NB, but that treatment effectively removed these cells in regularly budding GF *Hydra*. Dotted line indicates head and foot epithelial cell populations along the body column, and arrows highlight remaining foot-specific epithelial cells in head- and foot-less GF.NB. **D**. *In-situ* hybridization of the foot marker gene L-rhamnose-binding lectin CSL3-like (RBL) revealed it was exclusively expressed in the foot of regularly budding GF and GF.NB, but ectopic expression was visible in the distal body parts of GF.NB (arrow heads). Scale bar: 3mm. **E**. Activity of a foot-specific peroxidase was detected in the body column of GF.NB *Hydra* (arrow heads), confirming that localized ectopic terminally differentiated epithelial cells accumulated there. Scale bar: 3mm. **F**. Close-up of an ectopic foot region of the budding-inhibited GF.NB from E. Scale bar: 0.1mm.

### Epithelial stem cell genes that mark the differentiation into head-specific cells are inactivated in non-budding animals

The budding process is driven by tissue evagination and formation of a new head, that organises the formation of a new (secondary) axis (Bode, 2009). Hence, to understand the regulatory machinery controlling differentiation of epithelial stem cells in the context of the microbiota, we next used the scRNA-seq data to construct a complete trajectory from epithelial stem cells to terminally differentiated head cells. The trajectory analysis first of all showed that that there is no exclusively budding-specific trajectory: at the molecular level, the budding process is very similar to that of head formation (**Fig. 3A, C**). Examination of the gene expression profile in budding animals (WT, CV or budding GF) in both ectodermal and endodermal epithelial cells visualized the dynamics of gene expression during the differentiation process and identified the activation of numerous genes that appear to be involved in terminal differentiation into head epithelial cells. This gene activation program was drastically affected in non-budding animals in absence of bacteria (GF.NB). When comparing the profile of control animals (WT, CV, GF) with that of germfree non-budding animals (GF.NB), numerous genes were no longer expressed in the latter, both in ectodermal (**Fig. 3B**) and in endodermal cells (**Fig. 3D**). In total, 1115 inactivated genes were identified in GF.NB. Thus, associated with the non-budding phenotype, the epithelial cells seemed to be unable to execute their normal gene expression program in the absence of a microbiota. Among the genes lacking expression in GF.NB were many with known roles in stem cell differentiation and pattern formation in *Hydra*, including members of the WNT pathway. Genes encoding HyWnt1, HyWnt11, HyWnt16, and HyWnt 9/10c were expressed in both ectodermal and endodermal epithelial cells during head differentiation in the controls but not in GF.NB polyps (**Fig. 3B, D**). The genes HyWnt9/10a and HyWnt7 were specifically expressed in endodermal cells of the control polyps, while HyWnt3 was only expressed in their ectodermal cells. This indicates that our trajectory analysis indeed captured the gene expression dynamics during head formation in normal polyps and, in support of earlier findings (Hobmayer et al 2000; Holstein 2022, Holzem et al 2024), that the Wnt signaling cascade plays a key role in head-specific differentiation processes and also in budding.

**Fig. 3.**
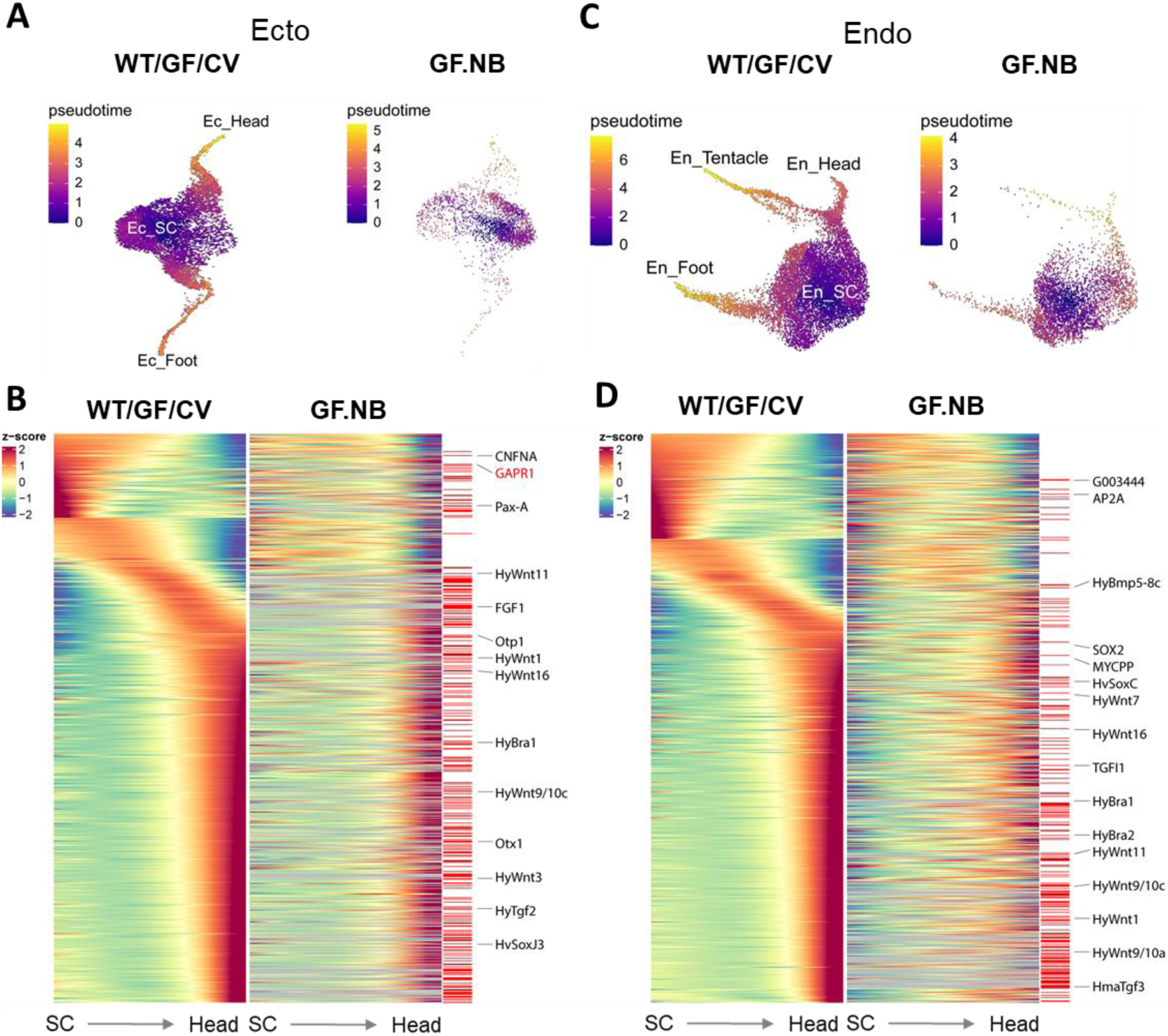
Expression of developmental key regulatory genes is disturbed in non-budding GF *Hydra.* Trajectory reconstruction for the ectodermal (**A**) and endodermal (**C**) epithelial cell lineages in regularly budding polyps (WT, GF, and CV) and in non-budding GF.NB. Corresponding gene expression dynamics is shown for ectoderm (**B**) and endoderm (**D**), collectively shown for regularly budding GF, WT, and CV, and separately for GF.NB. In these panels, the data are sorted from left to right for developmental cell rank (from stem cells, SC, to terminally differentiated head cells) and from top to bottom for peak expression over time. Genes initially expressed in the stem cells fade over time (top left section). The organized cell expression pattern of the budding animals is disturbed in GF.NB. All inactivated genes in GF.NB are marked by red bars next to the heat map and key genes are named. Scale bar: 0.5mm.

The trajectory analysis (**Fig. 3B, D**) further revealed highly relevant information on the timing of gene expression: even before expression of the classical pattern formation genes (e.g. BMP, WNT, HyBra), expression was detected of “early” genes that have so far not been associated with pattern formation processes in *Hydra*, and these were not expressed in the GF.NB animals. Interestingly, the expression dynamics of the two epithelial cell lineages revealed that in endodermal epithelial cells there were fewer of these “early” genes inactivated in GF.NB than in ectodermal cells (fewer red bars are visible in **Fig. 3B** than in **3D**). In addition, the large clusters of inactivated head-specific genes appeared much later in endodermal than in ectodermal epithelial cells. This is consistent with the fact that ectodermal epithelial cells, unlike endodermal epithelial cells, are in direct contact with the microbiota (**Fig. 1C**) and indicates that endodermal cells are affected by the absence of microbiota at a much later stage during their differentiation than ectodermal cells.

### Chromatin accessibility of epithelial stem cells are disturbed in non-budding animals

To explore whether the observed inactivation of epithelial cell genes in GF.NB involves global gene regulatory changes induced by the absence of microbes, we applied single-nucleus assay for transposase-accessible chromatin with sequencing (snATAC-Seq) to the same groups of *Hydra* used for scRNA-seq (**Fig. 2A**). Similarly, the three stem cell lineages of *Hydra* displayed distinct chromatin accessibility profiles while preserving the trajectories from stem cells to terminally differentiated cell types in each lineage (**Fig. 4A**). Though the overall chromatin accessibility profiles remained similar between groups, the number of nuclei of epithelial stem cells in GF.NB polyps differentiating towards head-specific cells decreased in both the ectodermal and endodermal cell lineages in comparison to controls (**Fig. 4B**). This is in line with the budding inhibition phenotype. Moreover, chromatin accessibility of both ectodermal (**Fig. 4C**) and endodermal (**Fig. S1**) epithelial cell specific genes of GF.NB polyps was lower compared to controls, suggesting a global regulatory remodeling in the absence of microbes. Further trajectory reconstruction based on the predicted gene activity from the chromatin profiles also revealed a temporal gene activation cascade following the ectodermal epithelial stem cell differentiation into head specific cells in normal budding polyps (**Fig. 4D**). Similar to the RNA-based trajectory (**Fig. 3B**), the gene activation cascade was also disturbed in GF.NB, and key development regulators including HyWnt7 and FGF1 were among the significantly suppressed genes (**Fig. 4D**). Among the suppressed genes appeared one interesting candidate, Golgi-Associated plant Pathogenesis Related protein 1 (GAPR1), which was one of the earliest affected genes in the SC→head trajectory of ectodermal epithelial cells in GF.NB (**Fig. 3B**).

**Fig. 4.**
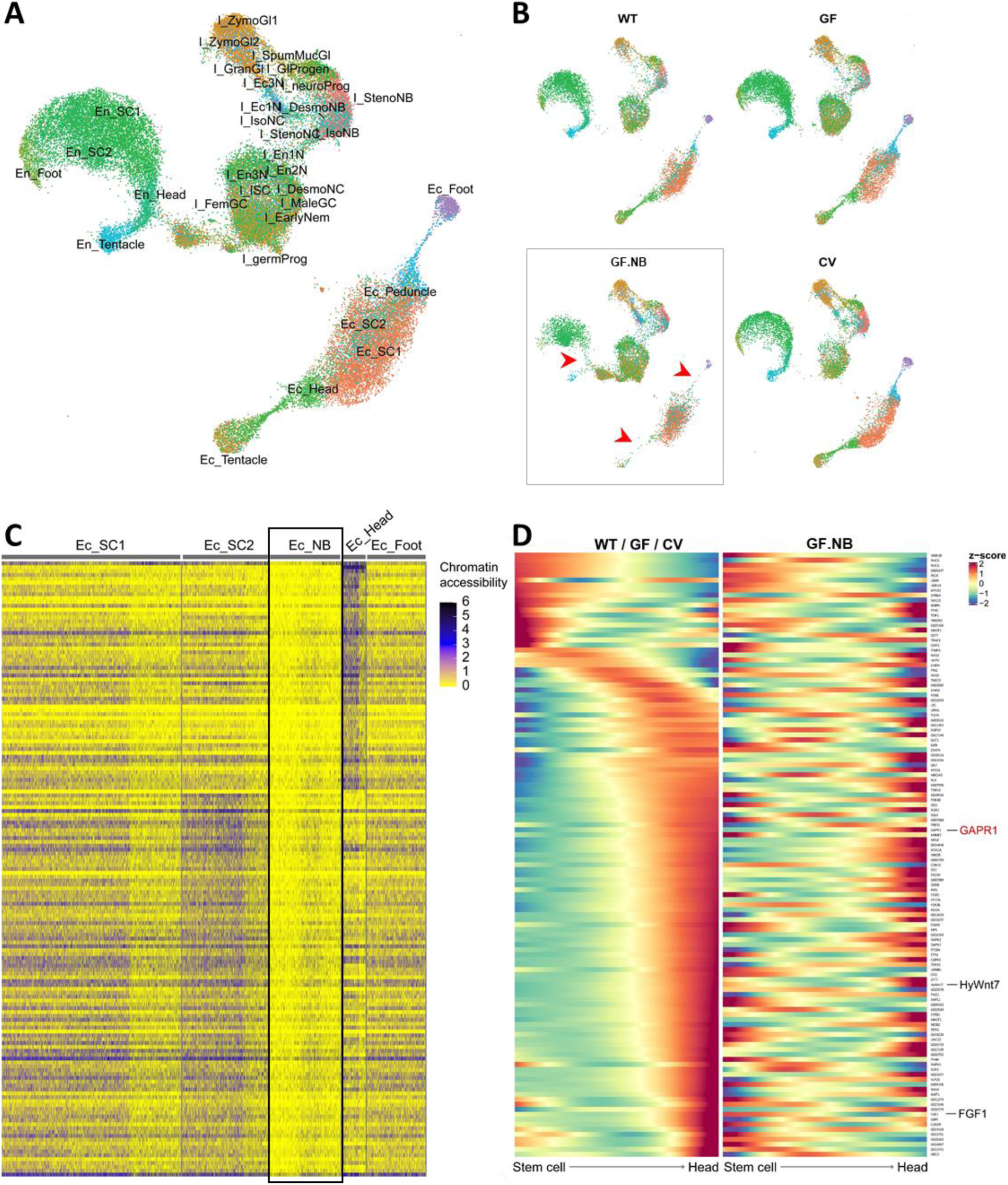
Microbiota interfere with epithelial cell chromatin openness in *Hydra*. **A**. UMAP representation of integrated single-nucleus chromatin accessibility profiles. **B**. Comparison of chromatin accessibility profiles between sample groups. Arrows indicate that in GF.NB polyps the number of ectodermal and endodermal stem cells terminally differentiating into head and foot cells is lower compared to controls. **C.** Ectodermal epithelial stem cells of GF.NB polyps showed distinct chromatin accessibility profile indicating low transcriptional activity. **D**. Comparative heatmap of chromatin accessibility predicted gene activity dynamics following the trajectory from ectodermal epithelial stem cells to terminally differentiated head epithelial cells, in regularly budding GF, WT, and CV *Hydra*, and in GF.NB.

### Candidate gene GAPR1 is causally involved in bud initiation

To validate the candidate budding genes, bulk RNAseq data were next produced from WT, CV, GF and GF.NB polyps, and from GF.NB polyps that had re-initiated budding spontaneously (GF.RB). The comparison identified 164 genes that were downregulated in GF.NB polyps but recovered in GF.RB (**Fig. 5A**). Inspired by the fact that ectodermal epithelial cells are responsible for the body shape of the animal (Wanek and Campbell 1982), by our observations that GF.NB animals have an obviously altered body shape compared to control animals (**Fig. 1**), and by the finding that ectodermal cells are more early affected by the absence of the microbiota than endodermal cells (**Fig. 3B, D**), a proof-of-principle analysis was performed with a selected single gene. From the many genes differentially expressed in ectodermal cells, GAPR1 was expressed early during the transition from stem cells to terminally differentiated head cells in control polyps but absent in GF.NB (**Fig. 3B**), and its expression was restored in CV polyps or once the budding was re-initiated (GF.RB). (**Fig. 5A**). By means of high-resolution in-situ hybridization, individual GAPR1 transcripts were visualized (**Fig. 5B**). In budding WT polyps, the GAPR1 transcripts were detected in ectodermal epithelial cells slightly below the newly forming bud (**Fig. 5B**, top series). A zoom showed their presence as fluorophor clusters corresponding to individual transcripts in ectoderm only. GAPR1 transcripts were never found in cells of the developing bud, but in detached buds, the transcripts became detectable in the lower body region upon maturation (data not shown). Interestingly, in well-fed polyps producing more than one bud, GAPR1 transcripts were also present in the ectodermal region just below the site where the next bud was being produced, opposite of the more mature bud that had already formed. In sharp contrast, no GAPR1 transcripts could be detected in GF.NB animals (**Fig. 5B**, lower series). This observation supports the findings from scRNAseq and bulk RNAseq data, and its observed early and localized expression suggests that GAPR1 may be causally involved in bud initiation, possibly by making ectodermal cells competent for budding.

**Fig. 5.**
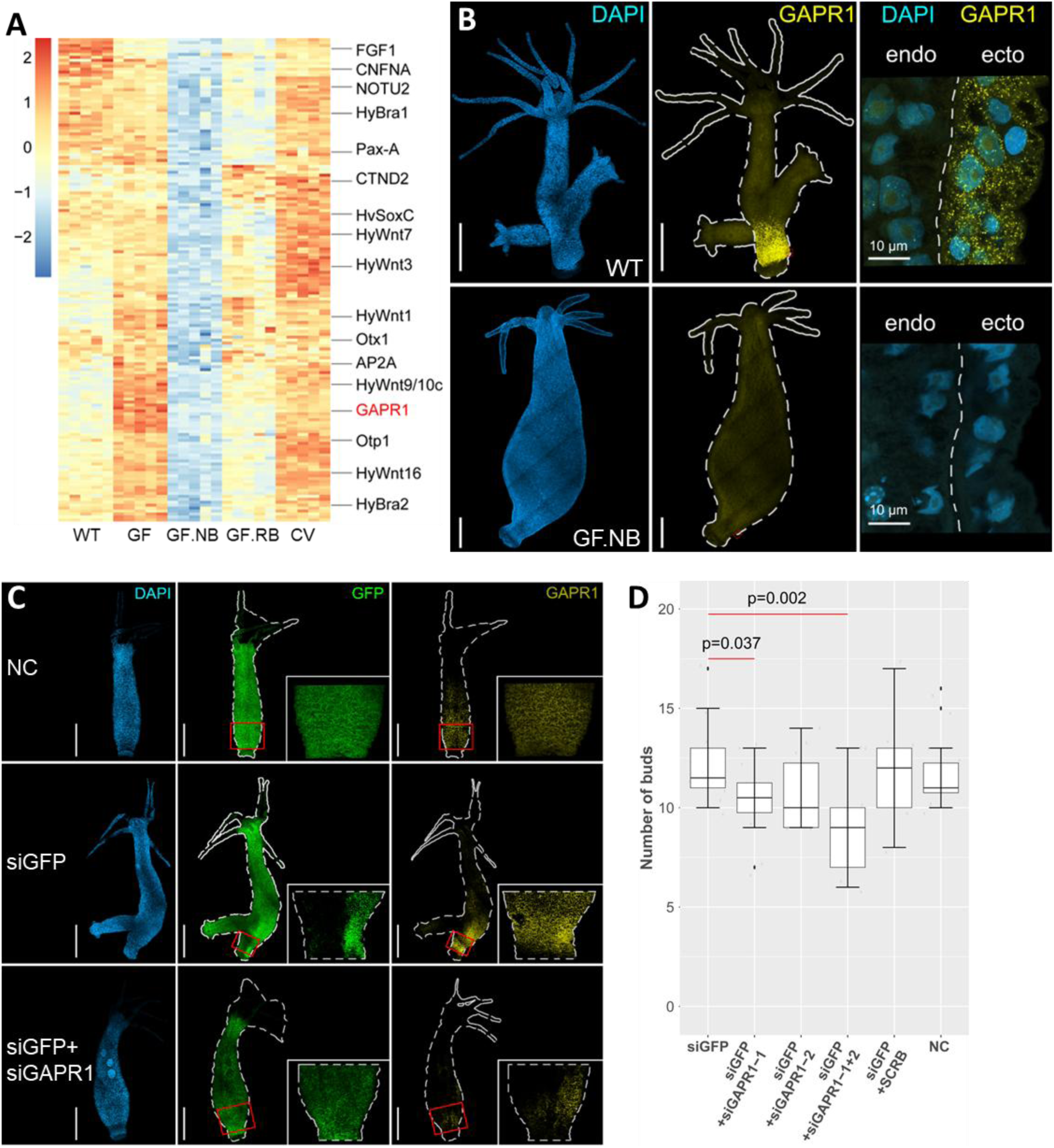
Microbiota interfere with key developmental pathways in *Hydra*. **A**. Heatmap showing the expression pattern of the inactivated genes in non-budding GF.NB, validated by independent bulk RNA-seq experiments. These genes were activated again when budding restarted in GF.RB. **B**. I*n situ* hybridization confirmed restricted expression of one of the early stem cell genes, GAPR1, in the peduncle region in WT budding polyps. No GAPR1 transcripts were detected in GF.NB polyps. **C.** I*n situ* hybridization confirmed targeted knock-down of GAPR1 (treated with siGAPFR1+SiGAPR2), as its expression was severely reduced 3 weeks after electroporation, while siGFP served as a positive control. The negative control (NC) received electroporation without any oligonucleotide. Scale bar: 2.5 mm. **D**. Knocking down GAPR1 by siRNA transfection with GAPR1-1 resulted in significantly reduced number of buds, with a stronger effect for siRNA silencing with GAPR1-1+GAPR1-2 in combination. The scrambled GAPRI siRNA had no effect. (n = 12 / group).

To determine whether the GAPR1 gene was causally involved in budding, we examined the effects of its transcriptional silencing on the budding process in GFP-expressing transgenic animals (Wittlieb et al 2006). For this, two different GAPR1-specific, short interfering RNA (siRNA) oligonucleotides (siGAPR1-1 and siGAPR1-2) were used alone and in combination, to induce gene silencing in intact *Hydra* polyps. Silencing of GFP by siGFP served as a positive control, and a control GAPR1 oligo with scrambled sequences (SCRB) was included. After their introduction into *Hydra* cells by electroporation, expression levels of GAPR1 transcripts were examined by in situ hybridization. **Fig. 5C** shows a drastic reduction of GAPR1 transcripts in polyps treated with GFP + GAPR1-1 + GAPR1-2, while electroporation with SCRB did not affect GAPR1 expression, indicating gene-specific silencing. Mock electroporated control animals (NC) expressed GAPR1 in the aboral region of the body column, as expected. Electroporation with the siGFP resulted in large areas without fluorescence, indicating the efficiency of the silencing (**Fig. 5C**). To test whether depletion of GAPR1 by RNAi also caused loss-of-function phenotypes, the budding process was monitored. Indeed, GAPR1 deficient polyps had a strongly reduced capacity for budding, in particular when electroporation was carried out with siGAPR1-1 and siGAPR1-2 in combination (**Fig. 5 D)**. The phenotypes varied between individuals, possibly due to variability in the fraction of effectively electroporated cells. Although the precise role of GAPR1 remains to be identified, the severe budding defects in the loss-of-function polyps indicate that this gene is functionally involved in bud initiation.

To summarize, by analyzing polyps that were significantly restricted in their budding behavior due to the absence of a microbiota, we identified a plethora of epithelial cell-specific genes that are expressed during bud formation, and not expressed when budding is halted in absence of a microbiota. A number of these code for factors that were already known to play a role in head differentiation and pattern formation, including budding, but we identified also a number of other genes that have not yet been associated with bud formation in *Hydra*. For a comprehensive understanding of pattern formation processes in *Hydra*, we consider both groups of genes to be equally important.

## Discussion

The holobiont *Hydra* provides a model to experimentally examine pattern formation in a simple multicellular animal, and demonstrates how this is affected by its microbiota. Our findings indicate that the ability of *Hydra* to reproduce asexually by budding is not solely attributable to host intrinsic properties; instead, the presence of colonizing bacteria has a significant external influence on whether the host animal can form buds or not, which is essential for the fitness of the asexually growing clone of *Hydra* polyps. Absence of bacteria temporarily halted budding, and had a major impact on chromatic accessibility and thus the epigenetic landscape. As a consequence of this and discovered by scRNA seq, many genes (including a number of yet uncharacterized genes) that are normally expressed in epithelial stem cell populations of budding WT were reversibly downregulated during a non-budding phase of GF animals. One of these genes is GAPR1; it is amongst the earliest active ectodermal genes in stem cells and is transcribed in WT even before the WNT genes are expressed. Its expression is localized near a newly-forming bud. A loss-of-function pilot experiment (**Fig. 5**) showed that its knockdown reduced budding capacity. How exactly the functionally important GAPR1 gene mechanistically affects the budding process remains to be investigated. In mammals, GAPR1 was proposed to act as negative regulator of autophagy (Shoji-Kawata et al 2013; Sheng et al 2019). In *Hydra*, previous studies have suggested that epithelial autophagy is required for maintaining epithelial self-renewal (Tomczyk et al 2020). The defects in budding caused by GAPR1 depletion (**Fig. 5**) support this view.

To interpret the data presented here, we return to Rahat and Dimentman (1982), who 40 years ago proposed a bacteria-produced budding factor in *Hydra*. Indeed, we observed a significant influence of the bacteria on the budding behavior of the polyps. However, the removal of the bacteria initially had little or no effect on budding behavior; the budding behavior of *Hydra* only decreased when they were cultivated without bacteria for a longer period of time. Even then, GF *Hydra* can still form buds, as we show in **Fig. 1**, though they form fewer buds and intermittently stop budding for a while. The animals therefore do not seem to lack a factor that has to be provided by symbiotic bacteria. We propose instead that during long-term absence of the microbiota, factors accumulate in or on the host that are inhibitory for budding behavior; in the intact holobiont these hypothetical factors would be continuously removed by the bacteria. Reintroducing bacteria in long-term sterile, non-budding animals resulted in an unusually large number of suddenly emerging buds, presumably resulting from the sudden and unregulated removal of this hypothetical inhibitory factor(s). This dependence of the asexual reproduction of the host on its bacteria may be the selective force that has established this ancient phylosymbiotic relationship.

Our work adds significantly to the previous observations of the profound impact of the microbiota on body shape and function of *Hydra*, while we concentrated on the effects on epithelial cells here. *Hydra*’s body size, for example, is determined not only by internal regulatory processes, but also by environmental factors, and also by the microbiota (Mortzfeld et al 2019, Taubenheim et al 2020). Like in other models (Beard and Blaser 2002), *Hydra* polyps increase in size when they lack a microbiota (**Fig. 1**) pointing to a conserved role of the microbiota in size determination. Our scRNA-seq and snATAC-seq data confirmed the importance of the ancient Wnt/b-catening signaling network in the development of both a head and of buds; in the absence of a microbiota, Wnt was downregulated in both ecto- and endodermal stem cells (**Fig. 3**). We also recently observed, that WNT/β-catenin signaling is more stable in presence of a microbiota (Taubenheim et al 2020). This work demonstrated that animals lacking a microbiota but exposed to a GSK3-3β inhibitor grew ectopic tentacles (Taubenheim et al 2020). Clearly, the microbiota of *Hydra* affects its developmental programs in multiple manners. It will be interesting to see what the function is of the less characterized genes that are expressed in presence of microbiota only during epithelial stem-cell maturation.

Whether this is a universal feature of animals is unclear. Developmental mechanisms evolved in the presence of colonizing microbes (Bosch and McFall-Ngai 2011; McFall-Ngai et al 2013; Bosch and McFall-Ngai 2021, Carrier and Bosch 2022). Removing or reducing microbial cells has substantial impacts on development, physiology and behavior of animals (Al-Asmakh and Zadjali 2015; Kennedy et al 2018; Argaw-Denboba et al 2024). In the absence of microbes, both in fish (Bates et al 2006) and mice (Reikvam et al 2011) cellular proliferation decreases. That animals are colonized by a specific and spatially organized microbiome with similarities over long evolutionary periods is recently recognized, but the ‘how and why’ provides an unresolved challenge in evolutionary biology and functional microbial ecology. The biochemical mechanisms by which bacteria influence the development of their hosts are still largely unknown. Given what we have learned from bacteria-free *Hydra*, we make the case that this model organism can not only provide insights into general principles of biological pattern formation, but also can pave the way to study the hidden impact of the microbiota on developmental pathways.

## Data avalability

All bulk and single-cell reads are available via the NCBI BioProject PRJNA1150130.

## Acknowledgements

This work was supported in part by grants from the Deutsche Forschungsgemeinschaft (DFG), the CRC 1182 “Origin and Function of Metaorganisms” (to TCGB) and the CRC 1461 “Neurotronics: Bio-Inspired Information Pathways” (to TCGB and AK). A.K. is supported by DFG grant KL3475/2-1. We are most grateful for the many discussions within the Bosch lab and also thank Christoph Kaleta for helpful discussions and Trudy Wassenaar for critically reading the manuscript. TCGB appreciates support from the Canadian Institute for Advanced Research, CIFAR. NGS analyses were carried out in cooperation with the CCGA at Kiel University. We thank the Central Microscopy Facility at Kiel University and Urska Repnik and Mark Bramkamp for their technical support. This research was also supported in part through high-performance computing resources available at the Kiel University Computing Centre.

## Materials and methods

### *Hydra* culture

All experiments were carried out using *Hydra vulgaris* strain AEP. Polyps were maintained at 18°C in sterile *Hydra* medium (S-HM) as previously described (Murillo-Rincon et al. 2017). All polyps were daily fed with GF *Artemia* nauplii, unless stated otherwise.

GF polyps were prepared as previously described (Franzenburg et al 2013). Briefly, wildtype polyps were incubated with a cocktail of antibiotics (50 µg/ml of Ampicillin, Neomycin, Rifampicin and Streptomycin, and 60 µg/ml Spectinomycin) for 14 days. The WT control received 0.01% DMSO (which was present in the cocktail) without antibiotics. On day 15, one group of GF polyps were incubated for 24 h with a supernatant of an overnight culture of homogenized WT *Hydra* (10 homogenized polyps per ml) to reintroduce a native microbiota, resulting in CV polyps. The sterility of GF polyps and of GF *Artemia* (that were hatched from decapsulated cysts with the same antibiotics cocktail used for generation of GF *Hydra*) was checked weekly by bacterial culture as described before (Murillo-Rincon et al 2017). In addition, PCR amplification of bacterial 16S rRNA genes was applied monthly to verify the absence of bacteria.

### Recording *Hydra* budding behavior

Individual polyps were cultured and monitored in sterile 24-well plates (#422.83.3922, Sarstedt), fed daily with GF *Artemia* and washed 14 h later. To synchronize the budding of individuals, an initial batch of 24 budding adults was randomly transferred into a fresh plate. The first newly detached bud of each animal was transferred into a second plate, and fed daily. The procedure was repeated by transferring a newly detached bud from each animal of this second plate into a third plate.

Budding dynamics of the polyps in the third plate was then checked twice daily, before feeding and after washing, for one month. The time of appearance and release of each bud was recorded, and newly detached buds were removed. No bud detachment lag was observed, therefore ambiguous bud detachment which may contribute to incorrect budding time records was absent.

### *Hydra* body size measurement

Body size was determined by measuring the total number of epithelial cells per polyp by flow cytometry as previously described (Mortzfeld et al 2019). For this, individual polyps were digested in 100 μL isotonic medium containing 75 U/ml Pronase E and released cells were analyzed by flow cytometry (FACSCalibu, BD Biosciences). Gating and acquisition parameters were determined by analyzing transgenic epithelial GFP-expressing polyps of the same *Hydra* strain (Mortzfeld et al 2019). Cells were counted and cell size was measured with FCSalyzer (0.9.18-alpha, https://sourceforge.net/projects/fcsalyzer/).

### Peroxidase activity staining

For detection of a foot-specific peroxidase (Hoffmeister & Schaller 1985), polyps were relaxed in 2% urethane for 2 min and fixed with 4% formaldehyde in PBS at 4 °C for 24 h. Following 24h incubation in 5% sucrose in PBS, samples were incubated with H_2_O_2_ in presence of 0.02% diaminobenzidine to reveal in situ expression as a brown coloring.

### Single-cell RNA-seq (scRNA-seq) analysis

Cell suspensions from 15 polyps per group were prepared as previously described (García-Castro et al 2021). Single-cell libraries, with three replicates per group, were prepared using the Chromium Single Cell 3’ Gene Expression kit (10X Genomics). Libraries passing in-house quality control checks were sequenced by NovaSeq 6000 S4 (Illumina). Raw reads were processed using Cell Ranger V7.2 (10X Genomics) following guidelines against the recently published chromosome-level genome assembly of *H*. *vulgaris* AEP (Cazet et al 2023). Quantification outputs were analyzed using Seurat V5.1.0 (Hao et al 2024), and pseudotime and trajectory reconstruction was performed using Monocle3 (Cao et al 2019). Data were plotted with the cell rank on the X-axis, sorted by their differentiation timing (pseudotime calculated by their expression profiles) and the gene rank on the Y-axis, sorted by their maximum expression timing. Genes sharing the same maximum expression timing were further sorted by their expression ranges i.e. how restricted their expression was across the sampled cells).

### Single-nucleus ATAC-seq (snATAC-seq) analysis

Nuclei suspensions from 10 liquid nitrogen snap frozen polyps per group were prepared using the Chromium Nuclei Isolation Kit (10X Genomics) following the user guide. Targeting 6000 nuclei per sample and two replicates per group, single-nucleus libraries were prepared using the Chromium Next GEM Single Cell ATAC Kit (10X Genomics). Libraries passing in-house quality checks were sequenced on two lanes of NovaSeq 6000 S1 (Illumina). Output raw reads were processed using Cell Ranger ATAC V2 (10X Genomics) following the default settings against the reference genome (Cazet et al 2023). Quantification outputs were analyzed using Signac (Stuart et al., 2021), and trajectory reconstruction was performed using Monocle3 (Cao et al 2019) as described previously.

### Bulk RNA-seq analysis

Total RNA from five replicates per treatment was extracted using PureLink RNA Mini Kit (#12183025, Invitrogen) and libraries were prepared following the Illumina TruSeq stranded mRNA (polyA enrichment) protocol. These were sequenced on 2 lanes of a NovaSeq 6000 S1 flow cell at the Competence Centre for Genomic Analysis (CCGA) Kiel using a 50 bp pair-end sequencing strategy. Quality control and preprocessing, including low-quality reads/ends trimming and adapter removal of the raw reads was performed using fastp (v0.20.0) (Chen 2023). Quality filtered reads were mapped to the new reference transcriptome using Bowtie2 (v1.2.3) (Langmead & Salzberg, 2012), and the aligned reads were sorted by Samtools (v1.9) (Danecek et al 2021) and further quantified using Salmon (v1.1.0) (Patro et al 2017). Quantification results were loaded into R (v4.4.1) (R Core Team, 2024) and analyzed with the DEseq2 (v1.26.0) (Love et al 2014) package in Rstudio (Rstudio Team, 2024).

### In-situ hybridization of GAPR1 transcripts

High-resolution in-situ hybridization of gene-specific transcripts was performed by hybridization chain reaction (Choi et al 2018), with modifications to the pre-treatment steps. Briefly, polyps were relaxed in urethane and fixed in formaldehyde as described above, and transferred into methanol for preservation. Upon use, the preparations were washed with 0.1% Triton-X 100 in PBS, digested in 10 μg/ml Proteinase K for 5 min, and fixed again with 4% formaldehyde for 30 min. The fixative was washed away as above, and the polyps were incubated in 5x SSCT at 70 °C for 20 min, prior to transfer into the hybridization buffer. All other procedures were according to the published method (Choi et al 2018).

### siRNA mediated gene knockdown

The protocol to knock down GAPR1 was performed following Reddy et al. (2019), using a transgenic line expressing GFP from the actin promoter. This allowed inclusion of a positive control with an siRNA oligo for GFP (siGFP), in addition to two oligos targeting different regions of the GAPR1 gene (siGAPR1-1, and siGAPR1-2). These were introduced, single or in combination, by electroporation. A control of GAPR1 scrambled (SCRB) sequences and a mock control electroporated without oligonucleotide were also included. The oligos were designed and ordered using the siRNA Design Tool from eurofins (https://eurofinsgenomics.eu/en/ecom/tools/sirna-design/). Twenty non-budding polyps were used per batch. Following the first introduction of the siRNA by electroporation the animals were allowed to recover and 4 days later the procedure was repeated. The resulting polyps were fed daily and their budding progression was recorded for three weeks.

---

**Fig. S1.**
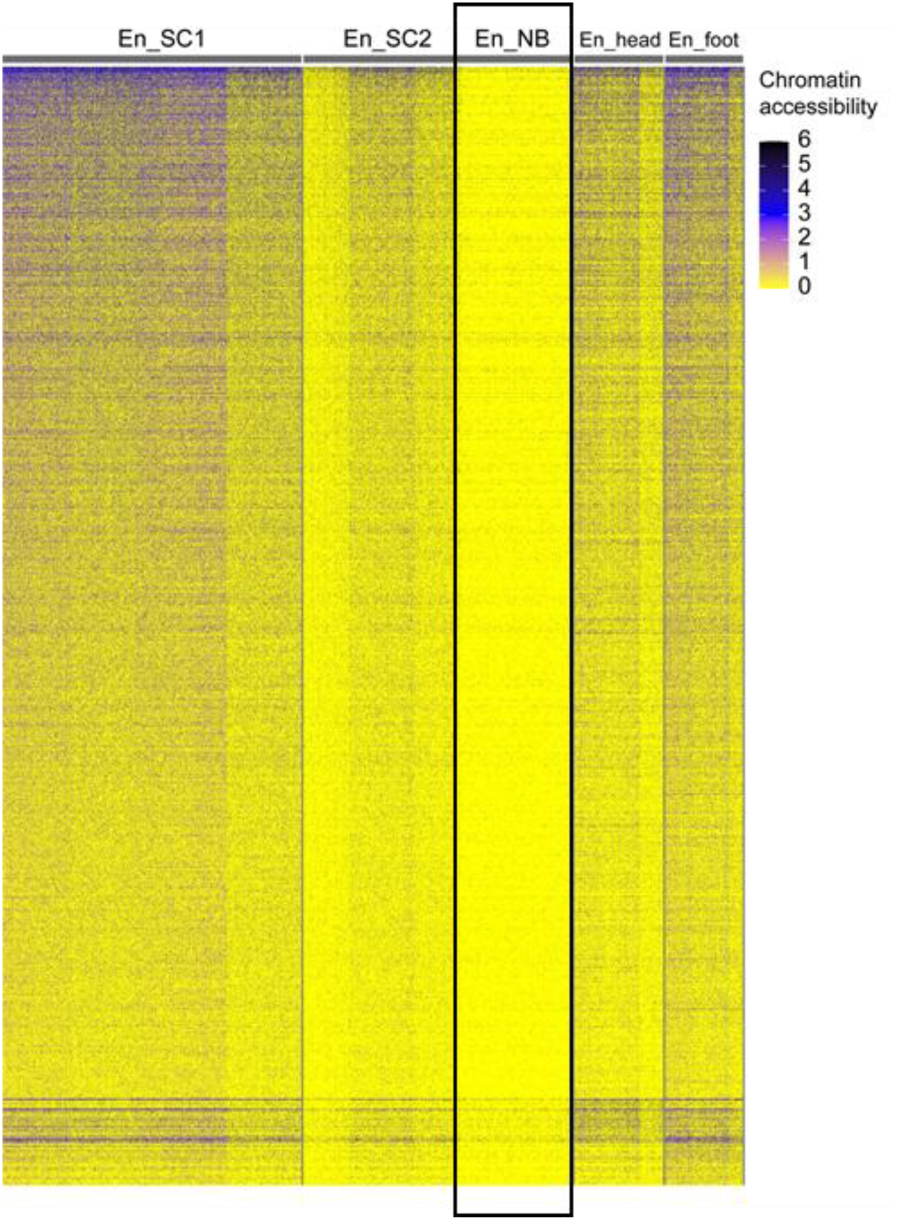
Microbiota interfere with endodermal epithelial cell chromatin openness in *Hydra*. Endodermal epithelial stem cells of GF.NB polyps showed distinct chromatin accessibility profile indicating low transcriptional activity.

